# Wide-field dynamic monitoring of immune cell trafficking in murine models of glioblastoma

**DOI:** 10.1101/220954

**Authors:** Elliott D. SoRelle, Derek Yecies, Orly Liba, F. Chris Bennett, Claus Moritz Graef, Rebecca Dutta, Siddhartha S. Mitra, Lydia-Marie Joubert, Samuel H. Cheshier, Gerald A. Grant, Adam de la Zerda

## Abstract

Leukocyte populations, especially tumor-associated macrophages (TAMs), are capable of mediating both anti- and pro-tumor processes and play significant roles in the tumor microenvironment. Moreover, TAMs have been shown to exert substantial influence on the efficacy of various cancer immunotherapy treatment strategies. Laboratory investigation into the behavior of TAMs has been limited by a lack of methods capable of resolving the *in vivo* distribution and dynamics of this cell population across wide fields of view. Recent studies have employed magnetic resonance imaging and intravital microscopy in conjunction with nanoparticle labeling methods to detect TAMs and observe their responses to therapeutic agents. Here we describe a novel method to enable high-resolution, wide-field, longitudinal imaging of leukocytes based on contrast-enhanced Speckle-Modulating Optical Coherence Tomography (SM-OCT), which substantially reduces imaging noise. We were able to specifically label TAMs and activated microglia *in vivo* with large gold nanorod contrast agents (LGNRs) in an orthotopic murine glioblastoma model. After labeling, we demonstrated near real-time tracking of leukocyte migration and distribution within the tumors. The intrinsic resolution, imaging depth, and sensitivity of this method may facilitate detailed studies of the fundamental behaviors of TAMs *in vivo*, including their intratumoral distribution heterogeneity and the roles they play in modulating cancer proliferation. In future studies, the method described herein may also provide the necessary means to characterize TAM responses to immunotherapeutic regimens in a range of solid tumors.

## Introduction

Tumor-associated macrophages (TAMs) constitute a critical cell population within cancer microenvironments, as indicated by the diverse roles TAMs play in mediating tumor cell proliferation,^1^ inflammation,^2^ and modifying tumor vasculature and extracellular matrix structure.^3^ This functional diversity arises in part from the presence of various TAM subsets across a phenotypic spectrum: M1-like polarized cells execute tumor-suppressing functions while M2-like polarized cells promote tumor survival and progression.^1^^,^^2^ To this end, therapeutic strategies relying on the induction of M1 polarization have been recently proposed for treatment of glioblastoma and other tumors.^4^^–^^6^ Furthermore, TAMs have been shown to respond to and influence the efficacy of radiation,^7^ immunotherapy,^5^^,^^8^ and other cancer treatment strategies.^4^^,^^9^^,^^10^

In this context, medical imaging technologies have proven indispensible to the exploration of TAM behaviors *in vivo* and their therapeutic potential and limitations. An especially useful strategy has been to exploit the robust phagocytic capacity of TAMs to accumulate nanoparticles that produce strong magnetic or fluorescent contrast, after which labeled cells can be detected *in vivo* using magnetic resonance imaging (MRI) or intravital microscopy (IVM)^4^^,^^9^^–^^12^ MRI and IVM respectively provide favorable imaging depth and sub-cellular resolution, however the physical detection limits of each method impede the simultaneous achievement of these properties. Optical Coherence Tomography (OCT)^13^ has been demonstrated as an alterative method for studying glioblastoma vasculature and infiltration at high-resolution,^14^^,^^15^ however this technique has not yet been adapted for detecting specific cell populations.

Our lab has recently reported contrast enhancement^16^ and image noise reduction^17^ methods that may facilitate the extension of OCT for studying TAMs and other tumor-resident leukocytes *in vivo*. In the current report, we explore the use of large gold nanorods (LGNRs)^18^ to label myeloid cells within the microenvironments of glioblastoma xenografts in mice. LGNR-labeled cells were subsequently detected *in vivo* with a modified OCT system, after which a battery of immunohistochemical methods was used to validate labeled cell populations in tumor and healthy brain tissues. Finally, we leverage high-speed three-dimensional acquisition to monitor the migration trajectories and speeds of individual labeled macrophages and activated microglia within tumors.

## Results

### High-resolution detection of LGNR retention in glioblastoma

Murine models of primary human glioblastoma were generated through the direct injection of U87MG cells into the brain parenchyma of *Foxn1^nu-/-nu^* mice, after which cranial windows were affixed to the exposed tissue to facilitate imaging. The crania of mice prepared in this fashion were imaged 7-8 days after tumor cell implantation using speckle-modulating OCT (SM-OCT), a modified implementation of OCT that improves resolution and minimizes speckle noise.^17^ Subsequent images were acquired following the intravenous administration of LGNR contrast agents as reported previously^16^ to achieve contrast enhancement within the tumor vasculature (**Figure 1a**). LGNRs exhibiting high shape monodispersity (102.0 ± 4.6 nm × 27.1 ± 1.7 nm) and spectral uniformity (λ_max_ = 825 nm, FWHM = 95 nm) were utilized because of their readily identifiable contrast when OCT images are reconstructed from multiple spectral subsets of the source bandwidth (**Figure 1b**).^16^^,^^19^^–^^21^

**Figure 1.**
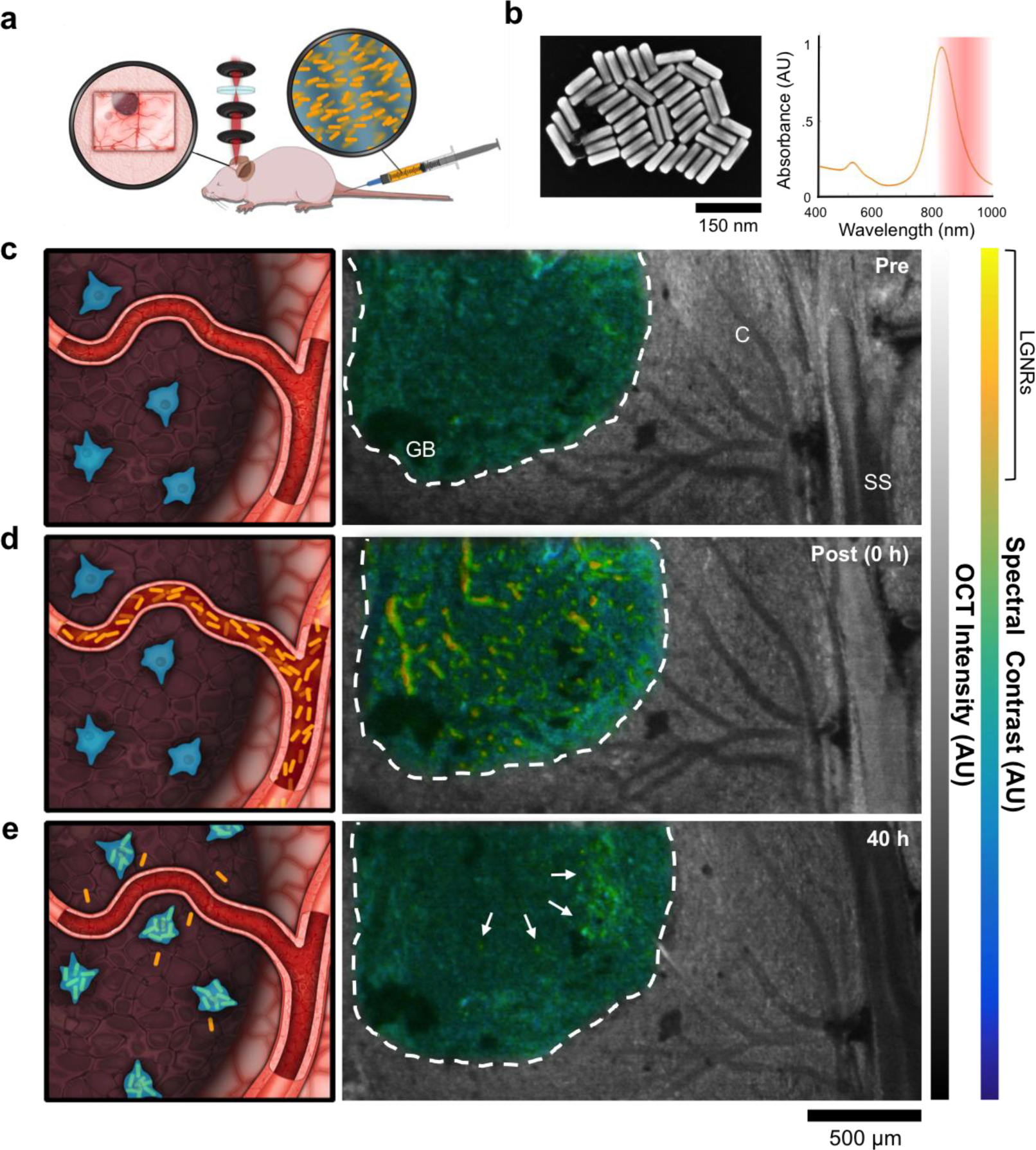
SM-OCT imaging of LGNR enhancement of glioblastoma vasculature and intratumoral retention. (a) Experimental setup. Mice bearing orthotopic glioblastoma xenografts were prepared with cranial windows and imaged with SM-OCT before and after injection of LGNRs. (b) Morphological and optical properties of LGNRs. (c) en face spectral SM-OCT image prior to LGNR injection. Glioblastoma (GB) margin is denoted with a white dashed line. Healthy cortex (C) and the sagittal sinus (SS) are also visible. (d,e) After injection, LGNRs reveal tumor vasculature (d) and exhibit prolonged retention within the tumor after circulation (e). Images indicate punctate distributions of LGNRs, possibly localized in cells.

Relative to surrounding brain parenchyma, models of invasive glioblastoma consist of dense cellular architecture, increased levels of associated host-derived immune cells, and abnormally tortuous vasculature.^22^ While these features were not readily apparent in pre-injection OCT images, glioblastoma could be distinguished from healthy cortical tissue by scattering intensity (**Figure 1c**). Administration of LGNRs by tail vein injection enabled detection of intratumoral blood vessels (**Figure 1d**). This signal persisted over hours, consistent with the measured half-life of LGNRs in circulation (t_1/2_ = 18 h).^16^ For each injected mouse, final images were collected at 40 h to assess the extent and distribution of LGNRs retained within glioblastoma resulting from the enhanced permeability and retention (EPR) effect.^[REF NEEDED]^ As a consequence of OCT’s innate high resolution, we were able to observe distinct, heterogeneous patterns of LGNR uptake that persisted long after the particles had cleared from circulation. For each mouse, we consistently noted numerous spectral puncta at 40 h within tumor tissue. To explain this observation, we hypothesized that punctate distributions of spectral signal within the tumor resulted from LGNR extravasation via EPR, followed by phagocytic clearance and accumulation within specific cellular subsets of the tumor microenvironment (**Figure 1e**). Informed by results we previously obtained for patterns of LGNR accumulation within liver and spleen tissue,^23^ we further predicted that the cells responsible for this uptake pattern likely originated from CNS myeloid populations.

### Assessment of intratumoral LGNR distribution

To corroborate and further explore these initial observations, we developed and implemented image segmentation methods to identify LGNRs from tissue using a combination of scattering intensity and spectral signal (hence called LGNR signal) calculated using dual-band Fourier methods. As expected, this segmentation revealed negligible LGNR signal within the tumor volume prior to injection. Segmentation results for images acquired immediately post-injection revealed the vascular architecture within the tumor. At 40 h, the LGNR signal is notably diffuse throughout the tumor and does not appear to correlate with the locations of blood vessels (**Figure 2a**). Methods were also developed to display the depth profiles of segmented LGNR signal, revealing the three-dimensional intratumoral distribution of particles (**Figure 2b**).

**Figure 2.**
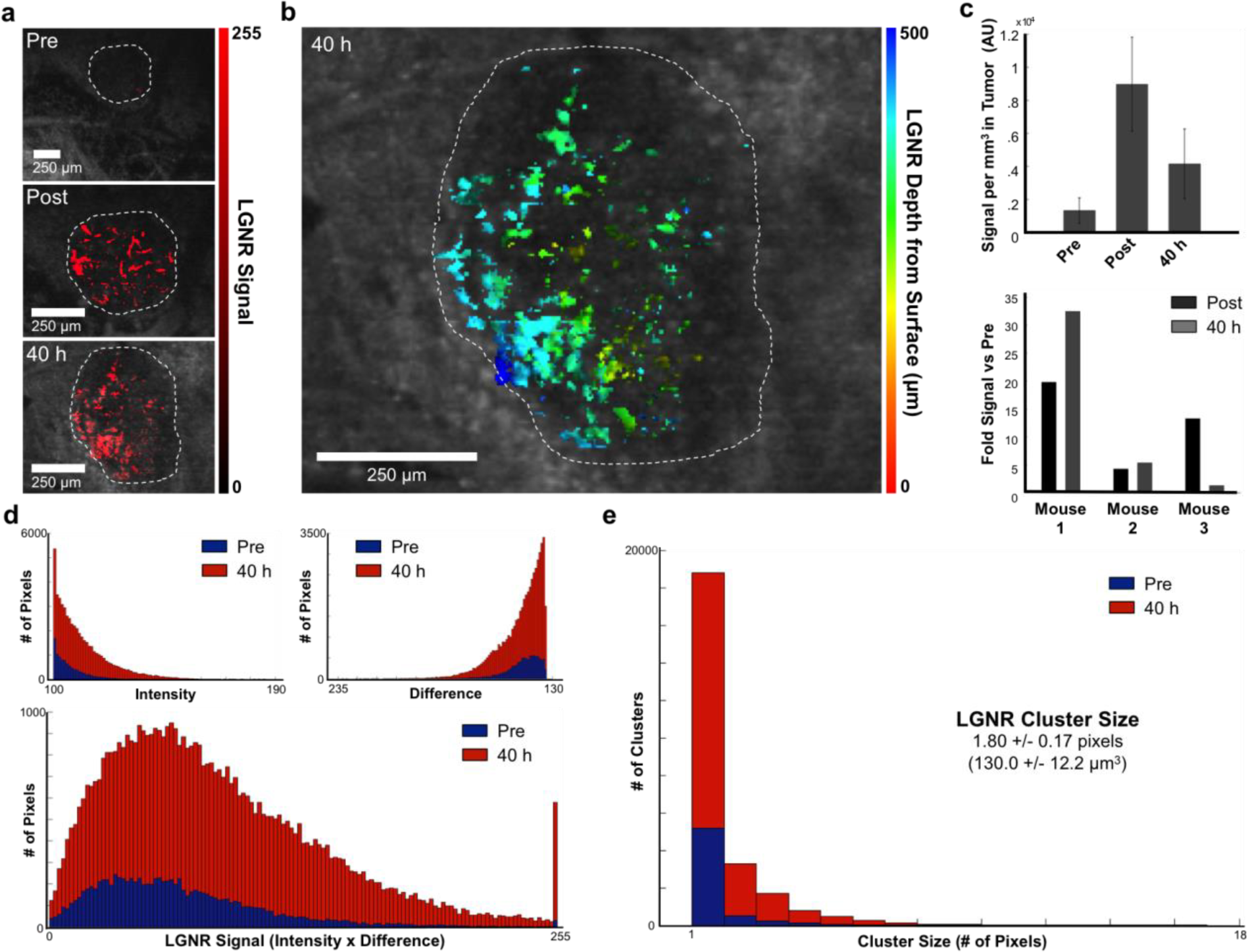
Characterization of LGNR distribution in glioblastoma. (a) SM-OCT images segmented based on spectral and intensity signals display LGNR localization in blood vessels immediately post-injection and distributed, often in clusters, throughout the tumor at 40 h. Tumor margins are outlined with white dashed lines. (b) 3D distribution of segmented LGNR clusters. LGNR cluster depth within the tumor is represented in color. In (a) and (b), segmented LGNRs are overlaid with a representative en face SM-OCT intensity image for anatomical context. (c) Average segmented signal values normalized by tumor volume for all time points and signal enhancement relative to pre-injection levels (n = 3 mice). (d) Representative distributions of pixels segmented by intensity, spectrality (difference signal), and LGNR signal (the scaled product of intensity and spectral signal) prior to injection and at 40 h post-injection. Segmentation threshold values can be tuned to change LGNR detection sensitivity and false positive rates (see Experimental Methods). (e) Distribution and average LGNR signal cluster size and approximation to cluster volume in cubic microns.

In general, we observed increased LGNR signal at both post-injection time points. However there is substantial variability across biological replicates, ostensibly arising from a number of variables including vascular density and permeability. This variability persists even when accounting for total tumor volume (**Figure 2c**, top panel). Expanding on this observation, there does not appear to be a consistent correlation among mice between the amount of signal observed in the vasculature immediately post-injection and the uptake observed at 40 h (**Figure 2c**, bottom panel). While we cannot definitively rule out differences that arise during model preparation, these results imply a notable degree of inter-tumor heterogeneity with regard to vascular density and the extent to which EPR can occur (i.e., vascular permeability is likely variable across tumors).

When measuring LGNR signal in this manner, it is critical to calibrate each segmentation parameter (see Experimental Methods) and implement the same parameters for images collected at all time points for a given biological replicate. Because LGNRs and tissue exhibit overlapping signal distributions, they cannot be distinguished using simple binary classifications, even when intensity and spectral signal thresholds are implemented. For this reason, we found it useful to compare the distributions of segmented voxels prior to LGNR injection and at 40 h to assess the extent of false positives (**Figure 2d**). As a note, segmentation thresholds for intensity and/or difference (spectral signal) parameters can be readily adjusted to minimize false positives, however this is necessarily done at the expense of LGNR detection sensitivity.

We further analyzed the size distribution of LGNR clusters, which we defined as contiguous regions of segmented voxels. When single-voxel clusters were considered, the mean LGNR cluster size was measured as 1.8 ± 0.2 voxels. Taking voxel dimensions into account, this corresponds to an average cluster volume of 130 ± 12.2 μm^3^ (**Figure 2e**). For consideration, this volume is roughly 24-42% of the mean cell body volume of healthy monocytes.^24^

### LGNR labeling of myeloid cells occurs *in vivo*

While the observed OCT results were suggestive of cell-mediated LGNR accumulation in tumors, a battery of *ex vivo* methods was required to rigorously test this interpretation. To this end, single cell suspensions were prepared from the excised brain tissues (healthy cortex and tumor) of LGNR-injected mice and analyzed using flow cytometry. Healthy brain and tumor samples from all mice consistently contained populations of CD11b^+^/CD45^low^ cells, which were interpreted as microglia.^25^^–^^28^ Relative to healthy brain samples, tumor samples from each mouse also contained substantial populations of CD11b^+^/CD45^high^ cells (**Figure 3a**). This expression profile, while non-comprehensive, is consistent with tumor-associated macrophages (TAMs),^25^^,^^26^^,^^29^ which are known to mediate both pro- and anti-tumorigenic phenotypes and constitute one manifestation of the immune compartment’s complex functions within the tumor microenvironment. Across the group of injected mice (n = 3), CD11b^+^/CD45^high^ cells constituted 5.2 ± 1.8% of all myeloid cells in healthy brain samples, compared with 24.6 ± 5.2% of myeloid-derived cells isolated from tumor samples (**Figure 3b**).

**Figure 3.**
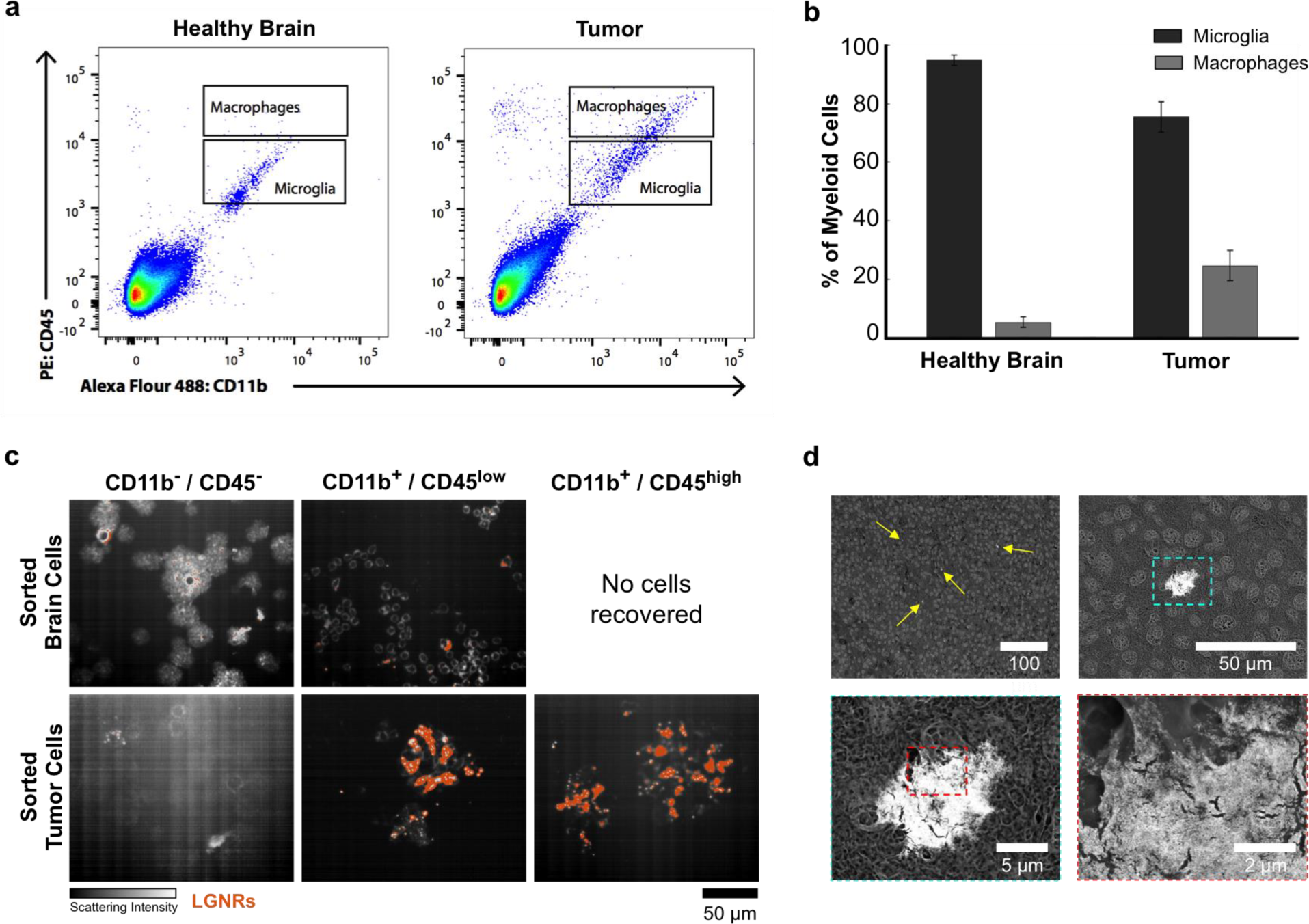
LGNR localization within leukocytes isolated from tumors. (a) Flow cytometry identification of putative microglia (CD45low/CD11b+) and macrophages (CD45high/CD11b+) in excised healthy and tumor tissues. (b) Microglia and macrophage quantification as percentages of total myeloid cell populations in healthy brain and tumor tissues (n = 3 mice). (c) HSM-AD detection of LGNRs (orange) in FACS-isolated cell populations. (d) SEM characterization of LGNR-labeled cells in histological tumor slices.

Once the cells were sorted, we sought to assess the presence of LGNRs as a function of cellular phenotype. To accomplish this, sorted cell suspensions were collected, concentrated, plated on standard microscope slides, and screened for the presence of LGNRs using hyperspectral microscopy (HSM-AD).^23^ Consistent with *in vivo* OCT results, cells isolated from healthy brain tissue (CD11b^−^/CD45^−^ and CD11b^+^/CD45^low^) displayed negligible presence of LGNRs. While CD11b^−^/CD45^−^ cells isolated from tumors did not contain LGNRs, both CD11b^+^/CD45^low^ and CD11b^+^/CD45^high^ cell populations exhibited LGNR accumulation (**Figure 3c**). These data indicated that some portion of LGNRs retained *in vivo* within glioblastoma were sequestered within myeloid cells exhibiting CD11b^+^/CD45^high^ and CD11b^+^/CD45^low^ expression profiles. CD45^low^ cells are most likely microglia, while CD45^high^ may represent a mixture of infiltrating TAMs and activated microglia. Moreover, the gold-derived contrast observed in scanning electron microscopy (SEM) images of paraffin-fixed tumor sections is also consistent with cell-mediated LGNR uptake (**Figure 3d**).

### Co-localization of LGNRs and myeloid markers in tissue *ex vivo*

We sought to further characterize the phenotypic profiles and distributions of LGNR-labeled cells through histological analysis of *ex vivo* brain and tumor tissue. Consistent with cytometry results, immunostaining indicated the presence of CD45^low^ cells within cortical tissue and CD45^high^ cells within the tumor. A subset of the cells within tumor tissue also stained positive for CD68^+^, a putative marker of phagocytically active macrophages (**Figure 4a**, top row). Immunostained slides were further analyzed using HSM-AD to detect the presence of LGNRs. Negligible LGNR signal was identified in sections of tumor-free cortex. By contrast, LGNRs were prevalent within tumor tissue and co-localized with CD45^+^ and CD68^+^ cells (**Figure 4a**, bottom row). In tumor tissue, more than 50% of the pixels that stained positively for CD68 were also positive for LGNRs. Using a separate method, 67.6% of pixel clusters (contiguous regions) that were CD68^+^ overlapped with LGNR+ regions (**Figure 4b**). These histological data further validated myeloid cell-mediated uptake as the origin of the intratumoral LGNR distributions observed during *in vivo* OCT imaging.

**Figure 4.**
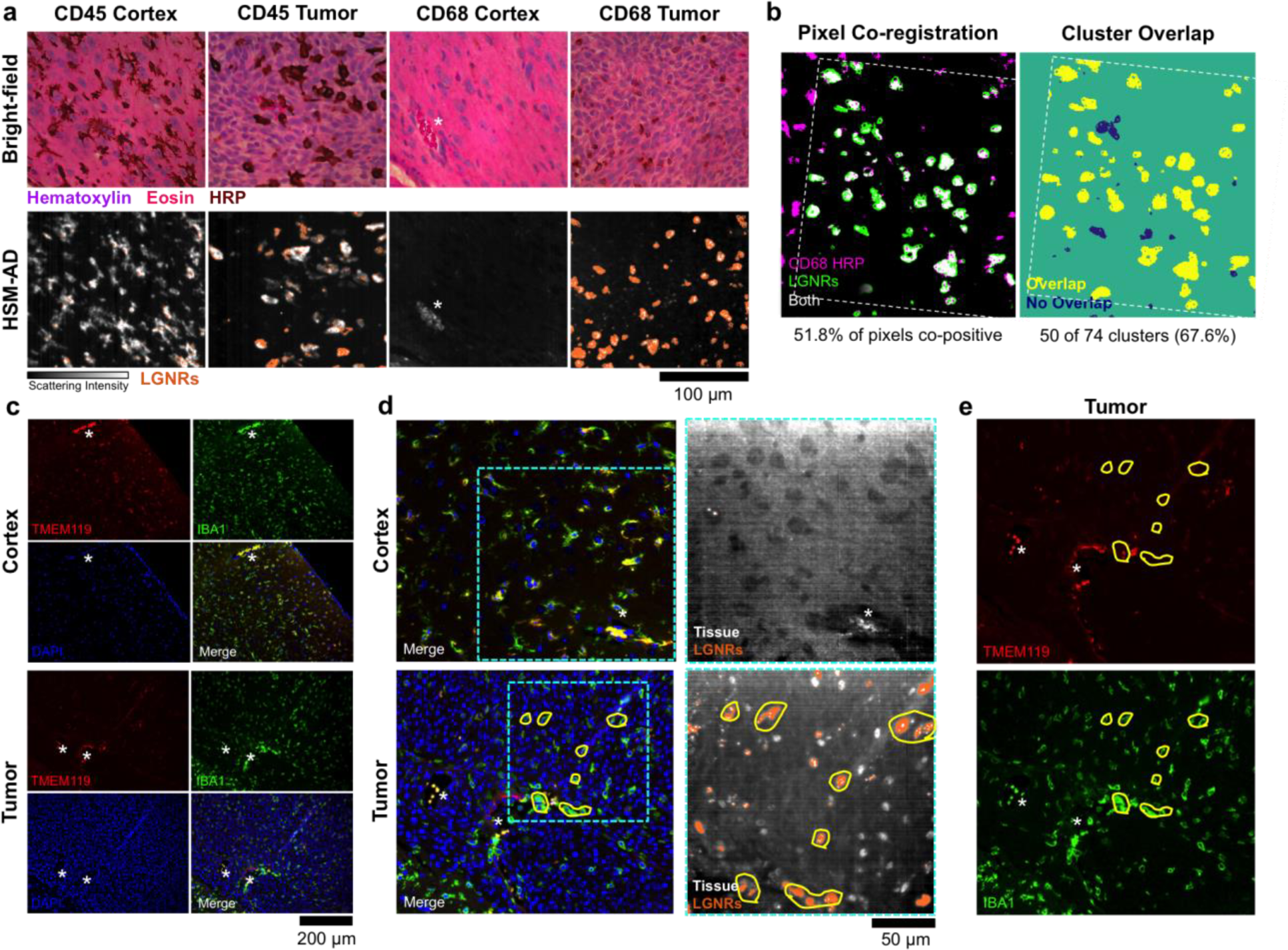
Immunohistochemical analysis of brain tissue sections and myeloid marker co-localization with LGNRs. (a) HRP staining of CD45+ and CD68+ cells in healthy cortex and glioblastoma (top row) and corresponding HSM-AD images revealing LGNR localization (orange, bottom row). (b) Image co-registration of segmented HRP staining and LGNRs reveal >50% co-localization of particles with cells expressing myeloid biomarkers. (c) Immunofluorescence staining indicates Iba1+/Tmem119+ cells in healthy cortex and Iba1+/Tmem119-cells in tumor tissue. (d) Comparison of fluorescence images to HSM-AD detection of LGNRs corroborates the absence of LGNRs in healthy brain tissue (top panel). In tumor tissue, LGNRs appear to co-localize strongly with Iba1+/Tmem119-cells. (e) Single-channel fluorescence images for Tmem119 (top panel) and Iba1 (bottom panel) annotated with regions of LGNR uptake (* denote blood vessels).

Additional tissue sections were analyzed to assess the origin of LGNR+ cells. Cortical tissue exhibited cells co-expressing Tmem119 (expressed by microglia)^30^^,^^31^ and IBA1 (expressed in microglia and macrophages).^32^^,^^33^ In tumor tissue, IBA1+ cells with faint or absent Tmem119 signal were abundant, however clearly Tmem119+ cells were infrequently observed (**Figure 4c**). Comparison of immunofluorescence and HSM-AD images indicated that LGNR+ cells in the tumor were IBA1+ with faint or absent Tmem119 signal (**Figure 4d,e**). These data suggest that LGNRs accumulate within tumor-associated macrophages, activated microglia, or both. Considered alongside results from cytometry, it appears likely that both cell types (which are closely related) actively accumulate LGNRs within the tumor stroma *in vivo*.

### Dynamic tracking of tumor-associated myeloid cells

Once labeled with LGNRs, intratumoral macrophages can be monitored over time. Starting 40 hours post-injection, 3D SM-OCT scans acquired every 15 minutes consistently indicate changes in the labeled leukocyte distributions within the tumor (**Figure 5a,b**). Superposition of segmented images acquired at adjacent time points reveals the migration trajectories of labeled cells over 90 minutes of observation (**Figure 5c**). Once segmented, trajectories of selected individual cells can be reconstructed over the entire tumor volume to enable comparisons of migration direction and velocity (**Figure 5d**). If volumetric scans are acquired at uniform scan intervals, changes in cell speeds can be approximated (**Figure 5e**), and quantified over time (**Figure 5f,g**). We measured an average speed of 3 μm/min for all identified labeled cells, which is consistent with previously reported macrophage migration speeds.^34^

**Figure 5.**
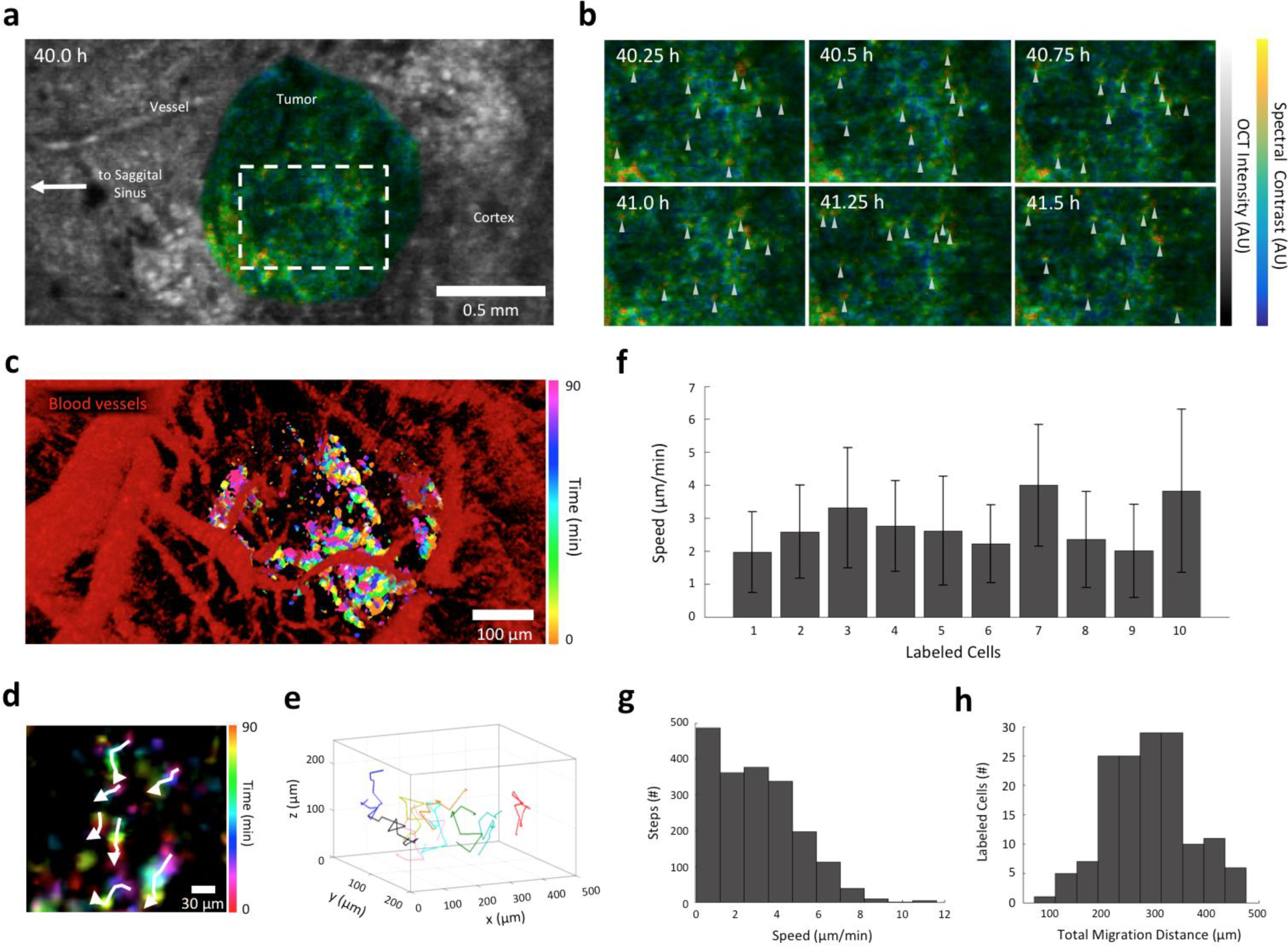
Dynamics of LGNR-labeled leukocytes in glioblastoma. (a) en face SM-OCT image of tumor (spectral, color scale) and surrounding brain parenchyma (intensity, grayscale) at 40 h post-injection. (b) Observed changes in the localization of LGNR-labeled cells (marked by arrowheads) within the tumor (images correspond to the dashed rectangle in (a)) sampled at 15 minute intervals. (c) Time-resolved monitoring of changes in the intratumoral distribution of labeled cells (color scale) in the context of tumor vasculature detected using OCT with speckle variance detection (red). (d,e) Mapping of labeled cell migration trajectories from 40 h post-injection to 41.5 h post-injection. In (e), 10 segmented cells were randomly selected for representation clarity. (f,g,h) Measurements of migration speeds (mean +/− s.e.m.) for selected individual cells (f), distribution of cell speeds calculated from adjacent imaging time points for all segmented cells (g), and total migration distance distribution for all segmented cells (h).

## Discussion

Efficient phagocytosis can be co-opted to densely label leukocytes with nanoparticles in cell cultures and in live animal tissues following systemic particle administration for the study of various diseases.^4^^,^^6^^,^^9^^,^^11^^,^^35^ Our results suggest a similar fate for intravenously-injected LGNRs. The absence of LGNRs within the brain parenchyma and presence within tumors indicates that particles extravasate through regions with a compromised blood brain barrier (BBB) and accumulate via the EPR effect. While EPR is known to be heterogeneous,^9^ this factor alone does not account for the patterns of discretely-localized LGNRs that we observe in tumors. Histological validation *ex vivo* strongly indicates that such patterns result from cellular sequestration of LGNRs. This data implies that, following extravasation, particles are actively collected by tumor-associated macrophages and microglia.

Tissue scattering and imaging noise pose substantial challenges to the complete and accurate detection of LGNR-labeled cells with OCT. While speckle modulation can significantly reduce coherent noise artifacts and improve LGNR detection in tissue, post-processing segmentation parameters must be chosen judiciously to minimize false positive detection. Processing images with low intensity and spectral signal thresholds improves labeled cell detection sensitivity at the expense of specificity and SNR. Conversely, application of strict thresholds may reduce false positives during cell segmentation while failing to identify cells that exhibit lower labeling density. These parameters are readily adjustable and should be optimized on an individual experiment basis.

Immunostaining results indicate that LGNR-labeled cells are tumor-associated macrophages. The presence of LGNRs in isolated CD11b^+^/CD45^low^ in addition to CD45^high^ populations suggests labeling of both tumor-resident microglia and infiltrating macrophages. It is known that reactive microglia induced by systemic inflammation or *in vitro* culture show reduced expression of many microglia signature genes including Tmem119.^30^^,^^31^^,^^36^ Based on the dramatically reduced intensity of Tmem119 staining in LGNR-labeled cells, it is likely that labeled microglia have adopted a tumor-reactive phenotype. While there is high correlation between LGNRs and various leukocyte markers it should be noted that not all leukocytes undergo labeling, and not all LGNRs end up within immune cells.

Contrast-enhanced SM-OCT offers powerful capabilities for tracking the distribution and single-cell migration dynamics of labeled leukocytes in tumors. In this capacity, the methods reported herein provide a new framework for exploring fundamental immunological processes that modulate cancer survival and proliferation. In light of recent studies, our method may be especially advantageous for monitoring macrophage responses to cell-mediated immunotherapies and more conventional cancer treatments. Previous work from our lab^37^ suggests that, in some cases, it may be possible to label immune cells with LGNRs *ex vivo* and track their uptake in tissue of interest following systemic administration. Future implementations of *ex vivo* labeling could expand the use of contrast-enhanced SM-OCT to study the dynamics of cell populations relevant to current cancer immunotherapy strategies.

## Acknowledgments

This work was funded in part by grants from the Claire Giannini Fund, the United States Air Force (FA9550-15-1-0007), the National Institutes of Health (NIH DP50D012179), the National Science Foundation (NSF 1438340), the Damon Runyon Cancer Research Foundation (DFS#06-13), the Susan G. Komen Breast Cancer Foundation (SAB15-00003), the Mary Kay Foundation (017-14), the Donald E. and Delia B. Baxter Foundation, the Skippy Frank Foundation, the Center for Cancer Nanotechnology Excellence and Translation (CCNE-T; NIH-NCI U54CA151459) and the Stanford Bio-X Interdisciplinary Initiative Program (IIP6-43). A.d.l.Z is a Chan Zuckerberg Biohub investigator and a Pew-Stewart Scholar for Cancer Research supported by The Pew Charitable Trusts and The Alexander and Margaret Stewart Trust. E.D.S. wishes to acknowledge funding from the Stanford Biophysics Program training grant (T32 GM-08294). O.L. is grateful for a Stanford Bowes Bio-X Graduate Fellowship. Research reported in this publication was supported by the Tashia and John Morgridge Endowed Postdoctoral Fellowship from the Child Health Research Institute at Lucille Packard Children’s Hospital as well as the National Institute of Neurological Disorders And Stroke of the National Institutes of Health under Award Number R25NS065741. The content is solely the responsibility of the authors and does not necessarily represent the official views of the National Institutes of Health. We also thank Christy Wilson PhD for training regarding the preparation of murine cranial windows, as well as Andrew Olson and the Stanford Neuroscience Microscopy Service (supported by NIH NS069375) for access to Imaris x64 software.

## Experimental Methods Contrast agents

LGNRs were synthesized, characterized, and adapted for biological use as described in prior work.^18^^,^^38^ Briefly, as-synthesized particles were coated with poly(4-styrenesulfonate) (PSS, MW=70 kDa) and thiolated methoxy-poly(ethylene glycol) (mPEG-SH, MW=5 kDa). Following surface functionalization and washing steps, LGNRs (λ_max_ = 825 nm) were prepared to 22.5 nM (OD 450 at peak λ) in water for intravenous injections as previously reported.^16^ Signal properties, detection sensitivity, and circulation half-life of these particles has been previously reported.^16^^,^^39^

### *In vivo* imaging system

All OCT images were acquired using a commercial OCT which was adapted to perform speckle-modulation (SM-OCT)^17^ for the removal of speckle and spectral-speckle noise. The commercial OCT was a spectral-domain High-Resolution system (Ganymede HR, ThorLabs, Newton, NJ), with a center wavelength of 900 nm and 200 nm bandwidth, which provide and axial resolution of 2 μm in water. The spectrometer of the OCT acquires 2048 samples for each A-scan at an adjustable rate of 1.1-30 kHz. Speckle modulation was implemented by inserting a rotating ground glass diffuser at a conjugate image plane. The main lens of the OCT provides a lateral resolution of 8 μm (FWHM) and depth of field (DOF) of 143 μm in water (LSM03-BB, ThorLabs, Newton, NJ). The ground glass diffuser (DG10-1500-B, ThorLabs, Newton, NJ) was rotated using an electromechanical mount (RSC-103, Pacific Laser Equipment, Santa Ana, CA). The optic axis was aligned with a radial offset relative to the center of the diffuser such that rotation ensured changing speckle patterns in the incident beam. The image plane was replicated by two lenses (LSM02-BB, ThorLabs, Newton, NJ) in a 4F configuration. Traditional OCT volumes were acquired in addition to SM-OCT volumes, by removing the diffuser from the optical axis.

### Cranial window implantation and orthotopic brain tumor animal models

The animal experiments were carried out in accordance with Stanford University Institutional Animal Care and Use Committee guidelines under APLAC protocols 26548 and 27499. Female athymic nude mice *(Foxn1^nu-/nu-^*, Charles River Labs) were anesthetized with ketamine/xylazine (100/10 mg/kg, intramuscular injection) and placed in a stereotactic frame. The scalp and underlying soft tissue over the parietal cortex were removed bilaterally. A drill was used to create a rectangular cranial window centered on the midsagittal suture that extended from the bregma to the lambdoid sutures. U87 human xenografts were implanted (10 μL of 1 × 10^6^ cells/ml cell suspension). Following implantation, a 5.5 mm × 7.5 mm × 3 mm glass cranial window was glued to the bone surrounding the cranial window.

### *In vivo* imaging protocol

Imaging was performed seven days after tumor-cell implantation. The mice were anesthetized with 2% isoflurane and placed in a stereotactic frame. The tail vein was cannulated and the mice were positioned for imaging. After initial (pre-injection) imaging, 250uL of OD 450 LGNRs (9x10^12^ LGNRs/mL) in PBS were injected intravenously. Mice were imaged immediately pre-injection, immediately post-injection, and at 40 hours post injection.

SM-OCT and OCT 3D volumes were acquired with 6 μm lateral spacing and 40 slow axis averages with an A-scan rate of 10 kHz for SM-OCT and 30 kHz for OCT. For SM-OCT acquisition, the diffuser was rotated at a tangential speed of ∼2 mm/s, which created non-correlated speckle patterns for each frame. Four 2.5 × 1 mm volumes were scanned and stitched to create a large field-of-view (5 × 2 mm) 3D volume from each mouse at every time point.

3D labeled cell tracking was performed by acquiring SM-OCT volumes at regular intervals (every 15 minutes for 90 minutes) and concatenating segmented volumes from each time point. At select time points, speckle variance images were collected with conventional OCT to produce blood vessel maps that were registered with segmented cell trajectories.

### SM-OCT signal and spectral processing

The structure of the tissue was obtained by conventional OCT reconstruction. SM-OCT provided frames with uncorrelated speckle patterns, which were intensity-averaged in linear-scale. All processing and analysis were performed with Matlab (MathWorks, Natick, MA).

The LGNRs can be detected owing to their unique spectral characteristics: a plasmonic spectral peak which provides enhanced scattering at one half of the OCT spectrum, compared to reduced scattering on the other half. When performing dual-band spectral analysis,^16^ LGNRs can be clearly distinguished from blood which does not contain LGNRs. However, similarly to conventional OCT signal, the spectral contrast also suffers from speckle noise, which has historically prohibited the detection of LGNRs in static tissue. Using SM-OCT, we were able to reduce the speckle noise of the spectral signal (spectral-speckles) and increase the visibility of LGNRs within the surrounding static tissue. In order to visualize both the OCT scattering intensity signal and he spectral contrast, we used a hue-saturation-value (HSV) representation of the image. The OCT signal was used to encode the value, which represents the brightness of the image. The spectral contrast encoded the hue (color) in the image. We used a Parula colormap to represent the spectral contrast. In this colormap, the tissue, which is spectrally neutral, appears bluish-green, whereas the LGNRs, which have a strong spectral contrast, appear in yellow-orange.

An additional spectral-noise reduction algorithm was applied in post-processing due to residual spectral-speckle noise (removing speckle noise entirely would require an infinite number of scans). In order to enhance the appearance of LGNRs and LGNR clusters, the de-noising algorithm relied on both the unique spectral contrast of the LGNRs and their increased scattering signal. We used a noise-reduction algorithm which resembles the traditional bilateral filter^40^ because it averages pixels based on their spatial proximity and radiometric similarity, however, our filter reduced noise from the spectral contrast and used the OCT scattering signal (non-spectral) as a metric for radiometric proximity. We used a fast-bilateral implementation.^41^ The result of this de-noising scheme maintains the spectral signal of the LGNRs while reducing spectral noise in the surrounding regions.

The depth inside the tissue is calculated by first detecting the surface of the tissue (under the cranial window) and then off-setting the depth of each pixel in the tissue by the distance of the surface of the tissue from the top of the image. Detecting the surface of the tissue is achieved by using a threshold to segment the tissue, which has a high OCT signal, from the region above the tissue, which has a low OCT signal. Errors in the detection of the tissue surface, for example due to noise, are eliminated by applying a two-dimensional median filter on the calculated depth of the surface of the tissue. In order to increase the speed of this calculation for very large volumes, we first subsample the tissue and detect its surface on a sparser grid instead of on every pixel. We then linearly interpolate the depth of the surface of the tissue on the full grid.

The LGNRs and LGNR clusters were segmented and quantified based on their LGNR-signal, which is defined as a combination of spectral contrast and OCT signal intensity (after SM-OCT averaging). First, thresholds for spectral contrast and OCT signal intensity are applied, meaning that a signal below the threshold is set to zero. Next, the remaining signal is linearized between 0 and 255. Last, the two metrics are multiplied to produce the LGNR-signal metric. This metric was used to quantify and segment pixels as LGNRs and it ensures that pixels that are segmented as LGNRs have strong spectral signal and high backscattering intensity. The parameters of the metrics: thresholds and slopes, can be tuned to according to the desired sensitivity and specificity. In this project, we tuned the segmentation parameters so that most of the visible LGNR clusters were segmented correctly. After segmenting the LGNR pixels we were able to differentiate between separate clusters by using a three-dimensional connected components algorithm (“bwconncomp”) with a connectivity parameter of 26. The size of each cluster, measured by the number of connected pixels, was obtained by the “area” property of each connected component (using “regionprops” function. The volume of each cluster in um^^^3 was obtained by translating the pixel size to μm. Manual frame-by-frame segmentation was used to identify tumor boundaries.

In order to track the LGNR clusters in time, the volumes acquired for macrophage-tracking were registered in three-dimensions to the volume of the first time-point using a rigid transformation which was calculated with the function “imregconfig.” Following registration, the segmented clusters were tracked in time by first finding the cluster centers (using the ‘Centroid’ property of “regionprops”) and then finding the closest cluster center in the subsequent time-points (using the function “dsearchn”). We calculated the motion-velocity of the clusters from their trajectories by taking into consideration the time between each acquisition, which was 15 minutes on average.

A color-coded time-laps image of the moving clusters was created by assigning different hues to each volume of segmented clusters acquired at each time-point and overlaying all of the segmented volumes by adding their pixel values.

The volume showing both the segmented LGNR clusters and the blood vessels was created by registering and overlaying the segmented volumes with a map of the blood-vessels, which was created by calculating the normalized speckle-variance^16^^,^^42^ from the conventional OCT volumes.

### Detection of LGNRs in hyperspectral dark-field images

We used a hyperspectral dark-field microscope (CytoViva, Auburn, AL) in order to detect LGNRs in tissue samples and in sorted cell populations from FACS. LGNRs are strong light-scatterers, which is why they appear with a high signal-to background in dark-field images. Additionally, they can be detected based on their unique scattering spectrum owing to their plasmonic peak. In order to detect and quantify the LGNRs in tissue samples, we used an automatic detection algorithm, which includes machine-learning classification, called hyperspectral microscopy with adaptive detection (HSM-AD).^23^^,^^43^ The detection algorithm was applied in post-processing using Matlab (MathWorks, Natick, MA). The pixels which are classified as having LGNRs are displayed in orange over the grayscale average-intensity image.

### Co-localization of LGNRs and immune biomarkers

Tissue slices were lightly H&E stained and immunostained for either CD45 (BD Pharmingen, 550566) or CD68 (Serotec, MCA1957) using horseradish peroxidase (HRP) methods (Histo-Tec, Hayward, CA). Stained tissues were imaged in bright field mode at various magnifications. The bright field images showing the HRP in dark-red were post-processed in order to segment the cells that expressed CD45 or CD68. Segmentation was performed on the blue channel of the RGB image, owing to the larger difference between the HRP and the surrounding stains in the blue channel compared to the other color channels or the gray-scaled image. The segmentation was implemented with Matlab using the function “graythresh,” which finds a global image threshold using Otsu’s method.^44^

The bright-field and the dark-field images were registered using Matlab’s control point selection tool (“cpselect”), a user interface that enables selection of control points in two related images. These control points are then used to calculate an affine transformation between the corresponding images. Next, the dark-field images were registered onto the bright field images to enable calculating the overlap between the segmented regions with HRP (from the bright field images) and regions with LGNRs (from the dark field images, after the HSM-AD algorithm). The co-localization was calculated both in straight-forward pixel-wise overlap and cluster overlap, in which the clusters were obtained with a connected-components algorithm (“bwconncomp”) applied to the segmented images.

### Cytometry and sorted cell analysis

Brains were removed and the tumor macroscopically dissected. Specimens were minced in RPMI, enzymatically dissociated at 37°C for 10 min with 5 mL TrypLE Express Enzyme (ThermoFisher Scientific: MA, USA) and 250 μg/mL DNAse1 (Worthington, USA). Following mechanical trituration, cells were filtered through 70 μm and 40 μm filters and subjected to a 0.9 M sucrose gradient centrifugation to dispose of dead cells and myelin. Upon ACK-lysis for removal of erythrocytes (Gibco Life Technologies, Carlsbad, CA, USA) cells were resuspended in FACS buffer containing HBSS, 1% bovine serum albumin, non-essential amino acids, sodium pyruvate, HEPES, EDTA, glutamine and antibiotic-antimycotic (Corning, NY, USA) and stained with the following antibodies:

CD45 PE (clone: 30-F11), CD11b Alexa Fluor^®^ 488 (clone: M1/70), CD31 Biotin (clone: 390), Ter119 Biotin (clone: TER-119), F4/80-Alexa Fluor^®^ 700 (all BioLegend, San Diego, CA, USA). Streptavidin Pacific Blue (Invitrogen, Eugene OR, USA). DAPI stain (Invitrogen, Eugene OR, USA) were used to exclude dead cells. Flow cytometric analysis and cell sorting was performed on the BD FACS aria II (Becton Dickinson). Appropriate fluorescence minus one controls were used to define the background gates.

Immediately after sorting, cells were spun down using a Cytospin™ 4 Cytocentrifuge. Cytoslides were fixed in 4% Para-formaldehyde (PFA) for 10 min and air dried. Cell samples sorted on the basis of CD45 and CD11b expression were then plated on standard microscope slides and imaged using the HSM-AD technique^23^ to assay the presence of LGNRs in each cell population.

### SEM sample preparation and imaging

LGNRs (5 μL of OD 1) were air-dried onto silicon wafer substrates and mounted onto 15 mm Aluminum stubs before visualization with a Zeiss Sigma Field-Emission SEM (Carl Zeiss Microscopy, Thornwood, NY) operated at 2-5 kV using InLens Secondary Electron (SE) and SE2 detection, and 5-7 kV using Backscattered Electron (BSE) detection. A combination of InLens and SE2 detection was used to characterize functionalized nanoparticle size and shape externally, while BSE detection highlighted internal rod-shaped particle size and shape, due to the relative high atomic number of the gold nanorods. Images were captured in TIFF using store resolution 2048 × 1536 pixels and a line averaging noise algorithm.

Tumor-containing brain tissue slices (5 μm thickness), which were previously formalin-fixed, embedded in paraffin wax and mounted onto glass slides, were deparaffinized with xylene and ethanol, rehydrated and prepared for SEM imaging. Tissue slices were contrasted with 1% aqueous OsO_4_ (30 min) and post-stained with 3% Uranyl Acetate (20 min), followed by 0.2% Reynolds’s Lead Citrate (2 min). Slices were rinsed well with ultrapure water (3x5 min) between individual contrasting steps, and vacuum-dried overnight in a desiccator before Carbon coating (Denton Benchtop Turbo, Moorestown, NJ) of slides to enhance conductivity. Brain slices were then similarly visualized with a Zeiss Sigma FESEM operated at 5-7 kV using BSE detection. Signal inversion resulted in micrographs with grey levels similar to traditional TEM images. Images were captured in TIFF using store resolution 2048 × 1536 pixels and a line averaging noise algorithm.

### Immunohistochemical analysis

FFPE sections were deparaffinized using xylene and rehydrated using ethanol/water washes. Antigen retrieval was performed by microwaving slides in boiling sodium citrate buffer (10 mM sodium citrate, 0.05% Tween, pH 6.0) for 1 minute, followed by continued incubation at room temperature (RT) for 15 minutes. Slides were blocked in PBS with 10% donkey serum and 0.5% Triton for 1 hour at RT, then stained with primary antibodies (Abcam ab209064 1 μg/mL, ab5076 1 to 500, Cell Signaling 3900S 1 μg/mL) in staining buffer (PBS with 1% donkey serum and 0.5% Triton) overnight at 4 degrees C. After PBS washes, florescent secondary antibodies were applied (Thermo A11055 and A21207, 1 to 500 each) in staining buffer for 2 hours at RT. Slides were mounted (Vectashield H1200) and imaged on a Zeiss AxioImager M1. Black levels for Tmem119 staining were established based on florescence observed using isotype control antibody. HSM-AD images of stained tissue sections (HRP and immunofluorescence) were also acquired to compare the co-localization of specific cell lineage biomarkers with LGNRs in the tissue.

